# A shared brain system forming confidence judgment across cognitive domains

**DOI:** 10.1101/2021.09.17.460809

**Authors:** Marion Rouault, Maël Lebreton, Mathias Pessiglione

**Affiliations:** Motivalion, Brain & Behavior (MBB) lab, Paris Brain Institute (ICM), Hôpital de la Pitié-Salpêtrière, Paris, France; Sorbonne University, Institut National de la Santé et de la Recherche Médicale (Inserm), Centre National de la Recherche Scientifique (CNRS), Paris, France; Laboratoire de Neurosciences Cognitives et Computationnelles, Inserm, Paris, France; Institut Jean Nicod, CNRS, Paris, France; Département d’Études Cognitives, École Normale Supérieure, Université Paris Sciences & Lettres (PSL University), Paris, France; Swiss Center for Affective Sciences (CISA), University of Geneva (UNIGE), Geneva, Switzerland; Neurology and Imaging of Cognition (LabNIC), Department of Basic Neurosciences, University of Geneva, Geneva, Switzerland; Paris School of Economics, Paris, France

**Keywords:** confidence, metacognition, time perception, semantic memory, belief judgment, functional magnetic resonance imaging, frontoparietal cortex, ventromedial prefrontal cortex

## Abstract

Confidence is typically defined as a subjective judgment about whether a decision is right. Decisions are based on sources of information that come from various cognitive domains and are processed in different brain systems. An unsettled question is whether the brain computes confidence in a similar manner whatever the domain or in a manner that would be idiosyncratic to each domain. To address this issue, human participants of both sexes performed two tasks probing confidence in decisions made about the same material (history and geography statements), but based on different cognitive processes: semantic memory for deciding whether the statement was true or false, and duration perception for deciding whether the statement display was long or short. At the behavioral level, we found that the same factors (difficulty, accuracy, response time and confidence in the preceding decision) predicted confidence judgments in both tasks. At the neural level, we observed using fMRI that confidence judgments in both tasks were associated to activity in the same brain regions: positively in the ventromedial prefrontal cortex and negatively in a prefronto-parietal network. Together, these findings suggest the existence of a shared brain system that generates confidence judgments in a similar manner across cognitive domains.

---

Most of our decisions are associated with a subjective estimate of their probability of being correct, known as confidence judgment (Fleming et al. 2012; Pouget et al. 2016). Humans can form confidence judgments across many levels of abstraction (Rouault et al. 2019) and multiple cognitive domains, such as perception, attention, memory and valuation – for reviews, see (Rouault, McWilliams, et al. 2018; Vaccaro and Fleming 2018). Confidence judgment is crucial for behavioral control, as it drives both arbitration between potential tasks – we tend to avoid situations in which confidence might be too low, and effort allocation to a given task – we tend to recruit more resources in situations where confidence is lower (Shenhav et al. 2013; Boureau et al. 2015). These considerations suggest that metacognition may be a domain-general process that enables ranking tasks on a unique confidence scale (de Gardelle and Mamassian 2014), just as options are ranked on a single valuation scale in economic decision theory (Levy and Glimcher 2012). The alternative hypothesis is that metacognition operates in a domain-specific fashion, with each task-related system generating its own confidence signal. Although fundamental for understanding the neural mechanisms underlying metacognition, the debate between domain-general and domain-specific views has yet to be settled.

A first approach to assessing domain-generality involves identifying the factors contributing to the formation of confidence judgment expressed by participants. Previous studies have shown that, for principled reasons, confidence is related to objective accuracy (Rouault and Fleming 2020), response time (Kiani et al. 2014) and difficulty (Kepecs et al. 2008). More accurate, faster and easier decisions are significantly associated with higher confidence, albeit to a different extent across studies (Sanders et al. 2016). Examining whether the relative weights of these factors in confidence judgments are similar across tasks can provide evidence for domain-generality. Nevertheless, similar contributors at the behavioral level can coexist with anatomical and functional separation between underlying metacognitive systems at the neural level (Baird et al. 2013; McCurdy et al. 2013), even if this possibility is less parsimonious than a single, shared metacognitive system.

A second approach to assessing domain-generality therefore consists in using functional neuroimaging to examine whether confidence judgments involve distinct or overlapping brain systems. Foundational work on the neural bases of confidence judgments has relied on perceptual decision-making as a model system. These early investigations have identified a frontoparietal network, centred on dorsal anterior cingulate and lateral prefrontal cortices (dACC and lPFC), in which activity is negatively associated with confidence level in both human and non-human primates (Kiani and Shadlen 2009; Hebart et al. 2015; Heereman et al. 2015; Bang and Fleming 2018). In contrast, activity in the ventromedial prefrontal cortex (vmPFC) has been positively related to confidence across perception, memory and valuation domains (De Martino et al. 2013; Lebreton et al. 2015; Gherman and Philiastides 2018; Rouault and Fleming 2020). However, it remains unknown whether commonalities across studies truly reflect domain-general metacognitive processes or simply shared features of cognitive tasks.

Addressing this issue requires collecting functional neuroimaging data while participants perform cognitive tasks that involve different high-level cognitive processes but using similar visual stimuli and motor responses (Fig. 1). A seminal attempt in that direction focused on the comparison of visual images, presented simultaneously in a perception task or after a delay in a memory task, and revealed a coexistence of both domain-general and domain-specific confidence signals (Morales et al. 2018). However, these two tasks shared at least partly overlapping cognitive processes, since they both involved the comparison of visual features. The conclusion about the domain-generality of metacognition was therefore limited to a subset of cognitive processes involving visual comparisons.

**Figure 1.**
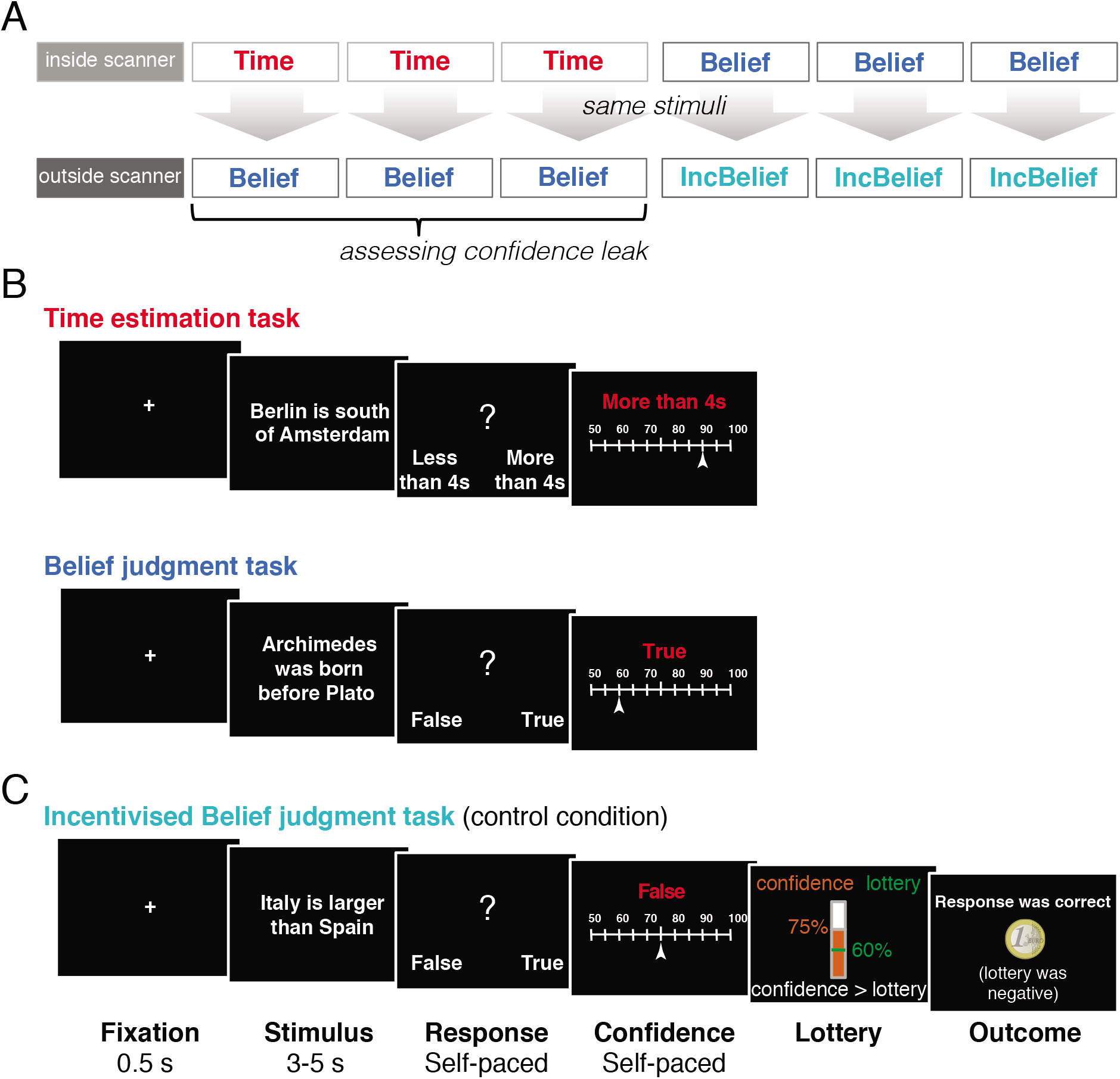
Experimental design. A) Schedule of task sessions. Inside the scanner, participants performed two experimental tasks, a time estimation task (“Time”, red) and a belief judgment task (“Belief”, blue). Afterwards, outside of the scanner, participants again performed a Belief task, now using the stimuli presented for the Time task in the scanner. Participants also performed an incentivised belief judgment task (behavioral-only) (“IncBelief”, green), using the same stimuli as those presented for the Belief task inside the scanner. B) Example of task trials. In all sessions, stimuli were general knowledge statements about history or geography facts. In the time estimation task, participants were asked to judge whether the statement was presented for more or less than four seconds on the screen. In the belief judgment task, participants were asked to judge whether the statement was true or false. In all tasks, participants were asked to rate their confidence in their response (reminded on the screen in red), on a scale from 50% (guessing) to 100% (sure correct). The incentivised belief judgment task (C) was similar to the belief judgment task, except that confidence ratings were now incentivised using a monetary bonus based on a lottery procedure – known as the Reservation or Matching Probability mechanism (see Methods). Confidence rating was compared to the winning probability of a random lottery, and the highest of these determined the bonus (+1€ if the response was correct or if the lottery was positive). Response accuracy and lottery outcome were provided on every trial.

Here, we intend to push further the notion of domain-generality with tasks that elicit radically different cognitive processes, while using the same stimuli and responses to avoid confounding fMRI contrasts. As stimuli, we presented short history or geography statements (e.g., ‘Kenya is larger than Gabon’). In the first task (time estimation), participants were asked to indicate whether the statement remained on screen for more or less than a reference duration (4 seconds). In the second task (belief judgment), participants were asked to indicate whether they believed that the statement was true or false. Thus, in the time estimation task, visual processing can be stopped at an early stage for just noticing when the stimulus is on and off screen without reading its content, whereas in the belief judgment task, the stimulus has to be read entirely so as to extract semantic information. The stimuli are therefore processed very differently in the two tasks, the information extracted from the stimulus being compared to an internal clock for time estimation and to semantic knowledge stored in memory for belief judgment. In both tasks however, participants were similarly asked to rate their confidence in their response.

A third approach to assessing domain-generality consists in testing for the presence of confidence leak between tasks, defined as an influence from a confidence judgment in a given task on a confidence judgment in another, unrelated task (Rahnev et al. 2015). To extend this concept, here we tested whether confidence leak may occur even from a task that is not overtly performed. In a previous study, we already demonstrated that neural confidence signals can be automatically generated, even if participants are not asked to provide a confidence rating (Lebreton et al. 2015). Here, we hypothesized that participants may not only read statements, but also spontaneously generate covert beliefs even during the time estimation task, despite such belief judgments not being required. We then reasoned that confidence in these putative covert beliefs may influence confidence in the overt response about time estimation. To search for behavioral and neural signatures of confidence leak, we collected belief and confidence judgments on the stimuli used for the time estimation task in a post-scanning session (Fig. 1A).

In summary, we developed a new paradigm involving two very different tasks, but the same confidence ratings, to assess domain-generality of metacognition via the three criteria described: similar weights of behavioral factors, overlapping brain activity and confidence leak across tasks.

## METHODS

### Participants

Twenty healthy human participants were recruited and scanned using functional magnetic resonance imaging in the neuroimaging center of the Paris Brain Institute (Cenir), within the Pitié Salpêtrière Hospital, Paris, France. Participants were screened for the following inclusion criteria: no history of neurological or psychiatric condition, no regular use of drugs or medication, and no contraindication to MRI (e.g. metallic implants). All participants provided written informed consent and the study was approved by the local Research Ethics Committee (Comité de Protection des Personnes, Inserm protocol C07-32). They were financially compensated for their participation by a fixed amount (75 euros), plus the money earned in the Incentivized Belief task (see below). The sample size was standard for neuroimaging experiments at the time of data collection (2009). One participant was removed for excessive head motion, leaving 20 participants for behavioral analyses and 19 participants for fMRI analyses (mean age=24, range 20-33 years old, 9m/10f). A power calculation done with G*Power toolbox (Faul et al., 2007), for the statistical tests used in data analyses with a power of 80% and a standard alpha threshold of 0.05 indicated that only moderate effects could be detected (unsigned Cohen *d*=0.58 for a significant one sample */*-test at the group level and unsigned coefficient ρ=0.50 for a significant correlation across participants).

### Experimental design

We built a new experimental protocol for assessing the properties of automaticity and generality of confidence judgments. Participants performed three experimental sessions inside the scanner and three outside of the scanner, of two different cognitive tasks with matched visuomotor properties (Fig. 1A). All tasks involved a 2-alternative forced choice followed by a confidence rating, all self-paced. The careful alignment of visual and motor requirements between tasks allowed us to eliminate low-level confounds in contrasts of neural activity aiming at isolating brain systems supporting the cognitive processes specifically involved in each task. The order of tasks was counterbalanced across participants.

#### Belief judgment task (“Belief”)

We were initially interested in selecting a task as best suited as possible for testing confidence automaticity. To facilitate confidence elicitation, we selected a task (i) for which the first-order response was likely to be generated (statements such as our stimuli being akin to common quiz games) and (ii) general knowledge to answer the statements is already present (or not) in memory, unlike metamemory paradigms in which participants are typically asked to encode and retrieve new material.

After a 0.5s fixation, participants were presented with a general knowledge statement and were asked to indicate whether the statement was true or false (Fig. 1B). The mapping between response keys and left/right presentation of the “true”/“false” responses was randomized across trials. Participants were then asked to report their confidence in their response on a rating scale from 50% to 100% correct (from chance level to perfectly sure by steps of 5%).

#### Time estimation task (“Time”)

After a 0.5s fixation, participants were presented with a general knowledge statement which stayed on the screen between 3s and 5s (as in the Belief task). Participants were asked to indicate whether the statement was presented for more or less than a reference duration of 4s. The mapping between response keys and left/right presentation of “more than 4s”/”less than 4s” was randomized across trials. Participants were then asked to rate their confidence in their response in identical conditions as for the Belief task (Fig. 1B).

#### Incentivised Belief judgment task (“IncBelief”)

The objective of this control condition was to compare confidence about identical stimuli across two different schedules of incentivisation for reporting confidence (Lebreton et al. 2018). We sought to take into account the consideration in the field of economy that probabilistic judgments or reports should be incentivised to be meaningful. Unlike subjective preferences which can be expressed in choices, it is less straightforward to assess subjective beliefs that are unobservable (Karni 2009; Schlag et al. 2015). This condition therefore allowed us to verify that participants respond to the best of their abilities, rather than producing somewhat random responses without a proper metacognitive evaluation.

It was identical to the Belief task except for the confidence report phase that relied on a Matching Probability procedure (also called Reservation Probability) (Hollard et al. 2016). First, participants rated their confidence on a scale as for the Belief task. Then, their rating was compared to a lottery of probability *p*. If their rating was superior to *p*, participants were rewarded 1 euro for a correct response and 0 euro for an incorrect response. In contrast if their rating was inferior to *p*, participants were rewarded 1 euro with probability *p* (and 0 euro with probability 1-*p*). Therefore, participants were incentivised to gamble on their own performance, taking responsibility for the outcome when sure about their response, and deferring to the random lottery when less sure.

#### Stimuli

Stimuli were general knowledge statements comprising 144 history statements, 144 geography statements and 36 blank statements (scrambled letters) (Fig. 1B). Stimuli were pseudorandomized across participants and across sessions. Critically, the same stimuli were presented during the three Time task sessions and the three Belief task sessions performed outside the scanner (Fig. 1A). Similarly, the same stimuli presented in the Belief task inside the scanner were presented in the IncBelief task outside of the scanner. Overall, each stimulus appeared once inside the scanner and once outside of the scanner.

### Behavioral analyses

#### Metacognitive ability

We analyzed accuracy (correct response rate), response time, and confidence rating for each task. We computed two metrics of metacognitive ability for each participant (for a review, see (Fleming and Lau 2014)). First, we estimated calibration, the difference between mean accuracy (objective) and mean confidence (subjective) in each task (Fig. S1). Second, we calculated discrimination, the difference between confidence ratings in correct and in incorrect decisions (Fig. S1), reflecting trial-by-trial metacognitive sensitivity. We note that the variability in difficulty level across trials for both tasks precluded the use of other metrics of metacognitive ability such as meta-*d*’. To compare these metrics between tasks, we conducted paired samples *t*-tests and Pearson correlation analyses. To further evaluate evidence in favor of the null hypothesis, we performed Bayesian paired samples *t*-tests with JASP version 0.8.1.2 using default prior values (zero-centered Cauchy distribution with a default scale of 0.707), and Bayesian correlation analyses with an a priori positive correlation, and reported Bayes factors (BF).

#### Contributors to confidence

Previous studies have shown that evidence available for the decision, determining the difficulty of the decision, influenced subjective confidence (Kiani et al. 2014). Note that difficulty here was defined as the opposite of the unsigned distance to the reference duration (4 seconds) for the Time task, and as the average accuracy measured in the other participants (all but the one tested) for the Belief Task. Despite it being an imperfect proxy due to a heterogeneous distribution of stimuli across participants and across conditions of incentivisation (Belief and IncBelief tasks), we found that it reliably captured decision evidence (Fig. 2), as it significantly predicted confidence ratings in all tasks, over and above other contributing factors. We additionally verified that no stimulus had an outlier signature (e.g. flooring or ceiling accuracy or confidence rating in most participants). Finally, as expected, we observed that accuracy and confidence correlated across all stimuli (ρ=.56, *p*<.0001). All possible factors susceptible to influence confidence judgment were included in a general linear model (GLM) regressed against confidence ratings to estimate their respective weights: accuracy, response time, difficulty, confidence in the preceding decision; along with five regressors of no interest: statement length (number of letters), statement duration, response (true or false), statement category (history or geography), and trial number. Regressors were z-scored to ensure commensurability of regression coefficients. A GLM was estimated for each participant, and beta coefficients were then assessed for statistical significance at the group level (using one sample *t*-tests against zero).

**Figure 2.**
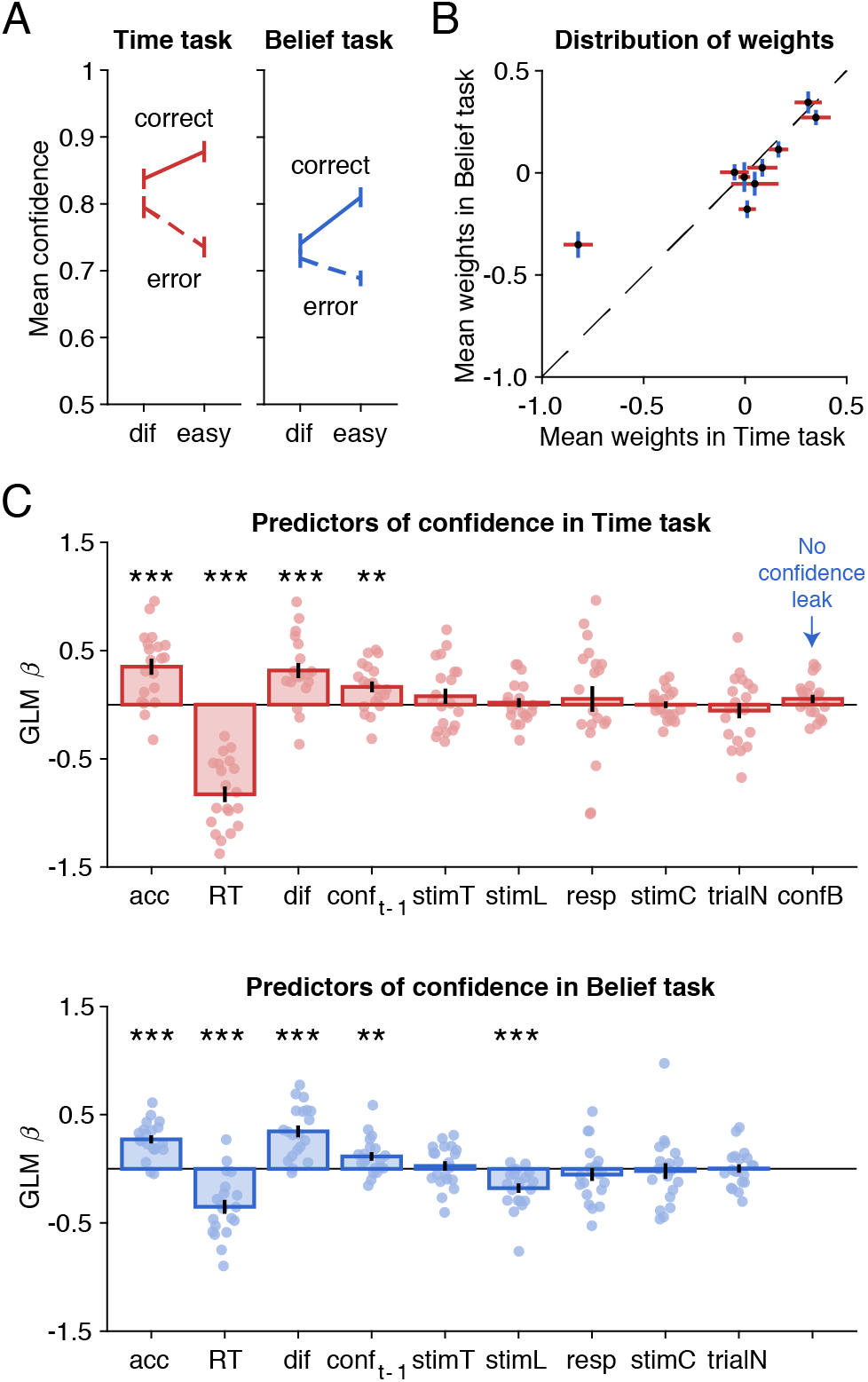
Psychometric properties of confidence judgments across tasks. A) Mean confidence in correct and error trials for easy and difficult decisions separately (median split of difficulty level). Circles and error bars indicate mean and SEM across participants (N=20). B) Correlation of weights associated with each factor between tasks (see C)). Error bars indicate mean and S.E.M. across participants (N=20) for each factor. C) Regression weights of factors predicting conf_t_ (confidence rating at trial t) in time estimation and belief judgment tasks. Accuracy (acc), response time (RT) and difficulty level (dif) all contributed to confidence. Regressors of no-interest were included for completeness: confidence at previous trial (conf_t-1_), stimulus presentation time (stimT), stimulus length (stimL), first-order response (resp, left or right), stimulus category (stimC, historic or geographic), and trial number (trialN). No evidence was found for a confidence leak (influence of confidence in belief judgment on confidence in time estimation (confB). Error bars indicate S.E.M. across participants (N=20). **p<.01, ***p<.001, two-tailed one-sample t-tests against zero.

Note that the correlation between weights in Fig. 2B is performed without the ‘confidence leak’ regressor in the GLM used to analyze confidence automaticity during the Time task, but with and without this regressor, the analysis provided virtually identical results.

### fMRI acquisition and preprocessing

We acquired T2*-weighted echo planar images (EPI) with blood oxygen-level dependent (BOLD) contrast on a 3.0 Tesla magnetic resonance scanner (Siemens Trio). We employed a tilted plane acquisition sequence designed to optimize functional sensitivity in the orbitofrontal cortex and medial temporal lobes (Deichmann et al. 2003; Weiskopf et al. 2006), with the following parameters: TR=2.0 s, 35 slices, 2 mm slice thickness, 1.5 mm interslice gap. T1-weighted structural images were acquired (1 mm isotropic, 176 slices), co-registered with the mean EPI, segmented and normalized to a standard T1 template, and averaged across participants to allow group-level anatomical localization. Imaging data were preprocessed and analyzed using SPM8 (www.fil.ion.ucl.ac.uk, Wellcome Trust Center for NeuroImaging, London, UK) implemented in Matlab. The first five volumes of each session were discarded to allow for T1 equilibration effects. Preprocessing consisted of spatial realignment, normalization using the transformation computed for the segmentation of structural images, and spatial smoothing using a Gaussian kernel with a full-width at half-maximum of 8 mm. Motion parameters from the realignment procedure were subsequently used as regressors of no interest in first-level analyses.

### fMRI analyses

Preprocessed fMRI time-series in each voxel of the whole brain were analyzed with a standard general linear model (GLM) at the participant-level and then tested for significance at the group-level. The main GLM (noted GLM1) included as regressors a boxcar function modeling the response time period, from stimulus onset to the first-order response (Fig. 1B), with a number of parametric modulations: response (true/false), accuracy (correct/incorrect), stimulus category (history or geography), statement presentation duration and statement length (number of letters). For both tasks, we modeled confidence as a separate Dirac function time-locked to the onset of the rating scale and parametrically modulated by confidence rating and rating time. Catch trials were assigned to a third separate regressor with statement presentation duration as a parametric modulator. Regressors of no interest included the six motion parameters calculated during realignment. The six scanning sessions (three Belief sessions, three Time sessions per participant) were modeled separately. All regressors were competing to explain variance (no orthogonalization). Regressors were z-scored to ensure comparability of regression coefficients and were then convolved with a canonical hemodynamic response function.

We created several contrast images showing confidence-related activity pooled across Belief and Time tasks, and confidence-related activity separately for Time and Belief trials (Supplementary Table 1). We extracted regions of interest (ROIs) surviving *p*<.05 familywise error (FWE) cluster-corrected for a cluster-defining threshold of *p*=.001 uncorrected. Regression coefficients were extracted from each ROI using a leave-one-out procedure. Significance was assessed at the group level (one sample *t*-tests against zero).

To examine generality of confidence-related activity, we extracted regression coefficients for confidence in the Time task from ROIs independently identified from confidence in the Belief task, and vice versa (Fig. 4) (uncorrected at the cluster level for positive correlation with confidence). For this analysis we used a cluster-defining threshold of *p*=.001 uncorrected (no FWE, because there was twice less data due to each task corresponding to half a scanning session). Finally, we computed two difference contrasts: confidence in the Belief task minus confidence in the Time task, and the other way around.

To investigate whether activity related to the first-order task was also domain-general, we contrasted activity at stimulus onset between Belief and Time tasks, and vice versa (Supplementary Table 2). Within each task separately, we further contrasted activity at stimulus onset between history and geography statements (Supplementary Table 3).

An alternative GLM (noted GLM2) was built to examine an automaticity hypothesis. For the Time task, we inserted three additional modulators of the boxcar function over stimulus presentation: response, accuracy and confidence related to the Belief task for the same statements. In a last variant (GLM3), we introduced these additional regressors as modulators of the stick function modeling the onset of the rating scale instead, but the conclusions were identical to those drawn from analyses using GLM2 (no neural correlate of confidence in belief judgment during time estimation).

In a final exploratory analysis, we investigated the relative contribution of neural activity in different regions of the domain-general network to behavioral confidence reports, separately for each task. Since the identification of ROIs in the domain-general network inevitably depends on statistical threshold, for this analysis we pooled together all negative ROIs obtained from the union contrast of both tasks. We estimated activity using one (stick) regressor per trial (deconvolution) and we then extracted and z-scored neural activity in positive and negative ROIs respectively, which then competed for variance in a bivariate regression analysis (Fig. 5).

## RESULTS

### Factors contributing to confidence judgment across tasks

Participants (N=20) were presented with history or geography statements (Fig. 1B) and were asked to judge whether the statement had been displayed for more or less than 4 seconds (time estimation task, thereafter “Time task”) or whether the statement was true or false (belief judgment task, thereafter “Belief task”). In both tasks, participants performed significantly above chance level, with an accuracy of 73.6% in the Time task (*t_19_*=16.7, *p*<1×10^-13^) and 67.2% in the Belief task (*t_19_*=18.1, *p*<1×10^-13^). There was a slight difference in accuracy between tasks (*t_19_*=-3.7, *p*=.0015), without a difference in accuracy between history and geography statements in the Belief task (*t_19_*=-.23, *p*=.82). As expected, confidence ratings were higher on correct than incorrect responses (Belief task, *t_19_*=11.5, *p*=5.2×10^-10^; Time task, *t_19_*=9.5, *p*=1.2× 10^-8^), denoting a degree of metacognitive sensitivity. We examined whether the sensitivity of confidence to difficulty level and to correct and error responses manifested similarly in the two tasks (Fig. 2A), through visualization of typical confidence patterns (Sanders et al. 2016).

Linear regressions revealed that the three expected factors had a significant influence on confidence ratings in the two tasks (Fig. 2C), namely: accuracy (both *p*<1.3×10^-4^), response time (RT, both *p*<2.6×10^-5^), and difficulty (both *p*<1.9× 10^-4^). Among other factors, confidence in the preceding trial response significantly contributed to confidence judgments in both tasks (both *p*<.0078), but the length of statements (number of letters) only affected belief judgment. We further verified that the results were robust to removing the predictors of no interest. We again found a significant contribution of accuracy (both *p*<7.7×10^-5^), response time (RT, both *p*<3.0×10^-5^), difficulty (both *p*<.0011), and confidence in the preceding trial response (both *p*<.0277) to confidence ratings in the two tasks. To assess the similarity in how the different factors impacted confidence rating, we computed the correlation of their regression weights between the two tasks (Fig. 2B). At the group level, this correlation was strongly significant (mean ρ=0.54, *t_19_*=8.61, *p*=5.57× 10^-8^).

To assess whether participants provided their best metacognitive judgment, they were asked to perform again the Belief task in a post-scanning session, where the accuracy of confidence ratings was incentivized using a lottery procedure (see Methods). In summary, to the best of our detection ability, the incentivization did not improve the accuracy of metacognitive judgments (Fig. S2 and Supplementary Results), suggesting that participants were already doing their best during the scanning sessions, in the absence of monetary incentives. Furthermore, the same significant factors were found to influence confidence judgments (Fig. S2), whether they were incentivized or not, and the weights of different possible factors were correlated between Belief and IncBelief tasks (mean ρ=0.60, *t_19_*=10.2, *p*=3.8×10^-9^). These results provide additional evidence for the robustness of the domain-general model, with the same factors contributing to confidence independently of the incentivization schedule.

We also examined several metrics of metacognitive ability, that specify how confidence judgment relates to accuracy: how close average confidence judgment is to average accuracy, and how well confidence judgment discriminates between correct and incorrect responses (see Methods). The inter-participant correlation of metacognitive abilities between tasks was around significance level (Fig. S1 and Supplementary Results), but this result should be taken with caution given the limited size of our sample (N=20).

### Confidence leak across tasks

We searched for a particularly demanding hallmark of confidence leak, which would manifest as an influence of confidence in belief judgment on confidence in time estimation about the same stimulus. For such a leak to occur, two cognitive steps are pre-required. First, the response (true/false) on the general knowledge statement should be spontaneously generated during time estimation. Second, the confidence in this belief judgment response should be automatically computed on top of it. If these pre-requirements are met, then confidence in belief judgment might influence confidence in time estimate. We tested this prediction by adding confidence ratings collected in the post-scanning session of the Belief task (using the same stimuli as for the Time task) in the GLM meant to explain confidence ratings in time estimates (see Methods and Fig. 2C). There was no significant influence of this additional regressor (*t_19_*=1.43, *p*=.169, BF=.548). Consistently, in a simpler regression analysis with confidence in belief judgments as the only predictor of confidence in time estimates, we again found no significant influence (*t_19_*=-.15, *p*=.88, BF=.233).

Concluding from this null result is problematic because the pre-required steps may not have occurred: participants may not even have read the sentence when doing time estimation, or generated a true/false judgment, or computed a confidence in this belief judgment. Going back one step, we replaced confidence rating by belief judgment (true/false) and/or response accuracy (correct/incorrect) in the GLM fitted to confidence rating on time estimates. There was again no significant influence of these two belief-related regressors on confidence in time estimation (all *p*>.12, all 1/3<BF<3). Thus, we found no evidence for confidence leak across tasks in the behavioral data. However, this does not preclude the possibility that the brain may still represent belief judgments and the associated confidence when performing time estimation, even if these representations do not affect confidence in time estimates expressed behaviorally. We investigate this possibility in the next sections.

### Overlapping brain confidence signals across tasks

To search for brain activity signaling confidence level, we fitted a GLM that included confidence rating as a parametric modulation, along with a number of regressors modeling other experimental factors (GLM1, see Methods). To assess whether there is a domain-general network for signaling confidence, we first examined confidence-related activity across tasks (union of contrasts) in a wholebrain analysis (Fig. 3A and Supplementary Table 1). Consistent with previous studies (De Martino et al. 2013; Lebreton et al. 2015), confidence ratings were positively reflected in the vmPFC (p_FWE_clu_=.0042). More precisely, the vmPFC cluster peaked at MNI [-2,50,-14], spanned across the left and right middle frontal gyri, Brodmann area 11, with most of the activity remaining anterior to the cingulate gyrus. We also observed a positive correlation with confidence in occipital regions, and a negative correlation in the contralateral occipital regions, but this is known as an artefact of the visual rating scale (more light enters the right brain when the eyes move to the right of the screen and vice-versa). We have used a vertical rating scale in subsequent studies (e.g., Abitbol et al., 2015), which eliminated the artefact in visual brain activity.

**Figure 3:**
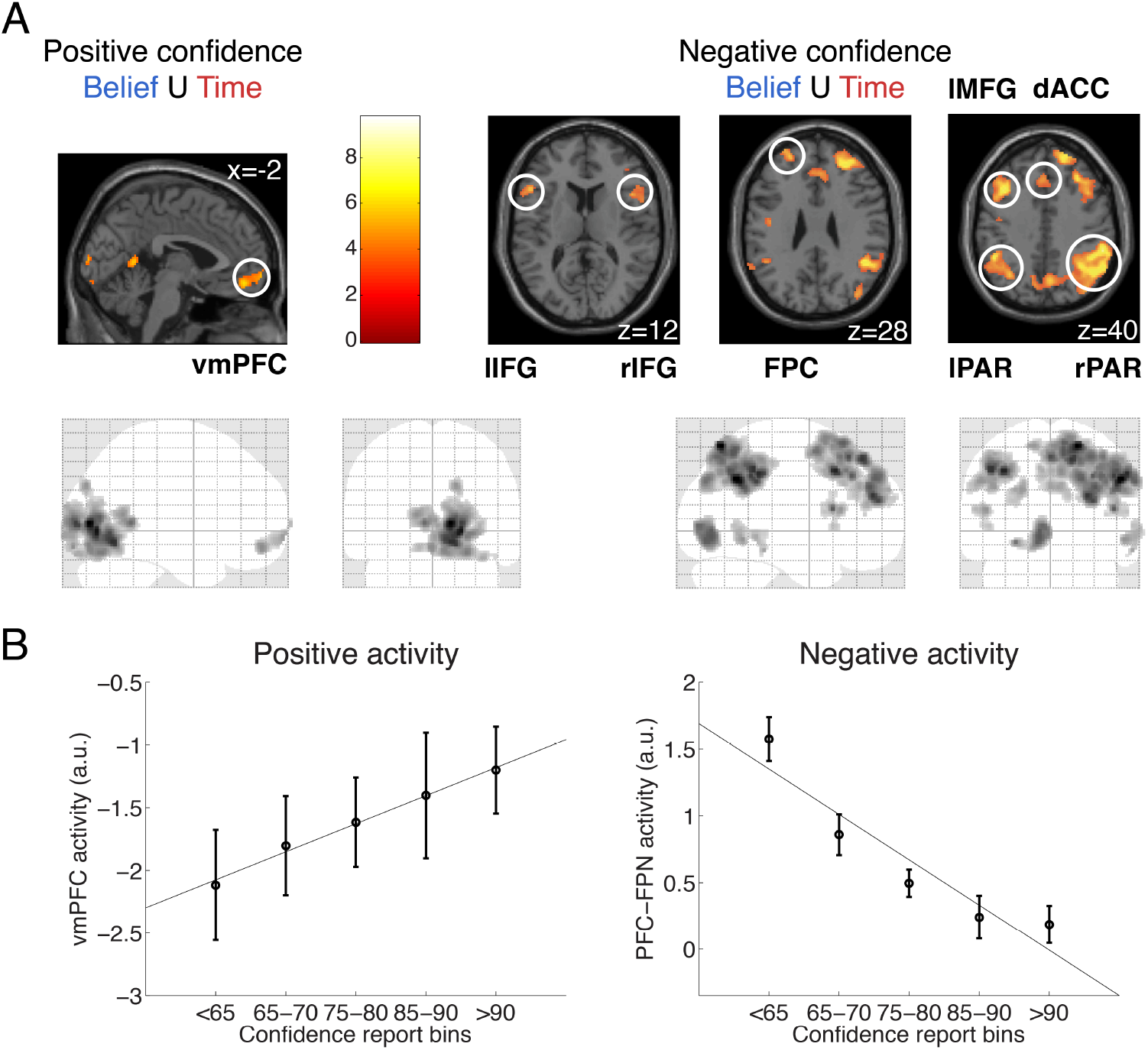
Domain-general confidence-related brain activity. A) Top panels: Statistical parametric maps show clusters in which activity is significantly associated with confidence rating (p<.05 after FWE cluster-wise correction for multiple comparisons, with a cluster-defining threshold of p=.001, uncorrected). Positive and negative associations are shown in left and right panels for the union contrasts (association with confidence observed across the two tasks). The color scale indicates the statistical T value in union maps. Numbers indicate x, y and z MNI coordinates of slices. Clusters are displayed at p=.001 uncorrected, k>60. Bottom panels: Glass brain maps corresponding to the two union contrasts (k>140 for display purposes). Abbreviations: vmPFC (ventromedial prefrontal cortex), IFG (inferior frontal gyrus), FPC (frontopolar cortex), dACC (dorsal anterior cingulate cortex and supplementary motor area), MFG (middle frontal gyrus), PAR (parietal cortex), l (left) and r (right). See also Supplementary Table 1. B) Positive activity in the vmPFC (left panel) and negative activity in the prefronto-parietal network (right panel) for bins corresponding to portions of the confidence scale, pooled across tasks. Error bars represent SEM across participants (N=19).

In addition, we identified a large prefronto-parietal network in which activity was negatively associated with confidence across tasks (Supplementary Table 1). The prefrontal clusters included a large region along the medial wall, including the dACC and supplementary motor area, and extending laterally (thereafter, “dACC”, p_FWE_peak_=.0188). We also found significant clusters in lateral regions in the inferior frontal gyrus (left, “lIFG”: p_FWE_clu_=.0205; right, “rIFG”: p_FWE_clu_=.0070), middle/inferior frontal gyrus (“lMFG”: p_FWE_clu_=5.6×10^-7^), and more rostral regions in the frontopolar cortex (“FPC”, p_FWE_clu_=.0039). The parietal clusters were located in the precuneus, superior and inferior parietal lobules (left, “lPAR”: p_FWE_clu_=7.5×10^-12^; right, rPAR: p_FWE_peak_<.0013) (Fig. 3A). In contrast to positively correlated activity, which was focal in the vmPFC, negatively correlated activity was much more widespread across the brain and more strongly associated with confidence (Supplementary Table 1). We also verified that positive and negative correlations were not restricted to high and low confidence ranges, by showing activity (pooled across tasks) in bins corresponding to adjacent portions of the confidence rating scale (Fig. 3B).

To confirm that the same voxels were signaling confidence in both belief judgments and time estimates, we conducted a cross-over analysis, extracting confidence weights on activity recorded in one task, within regions of interest (ROI) independently defined using the other task (as group-level significant confidence-related clusters) (Fig. 4). Regarding positive confidence signals, we found significant weights on activity during the Belief task within the unique ROI identified from the Time task in the vmPFC (*t_18_*=2.57, *p*=.0193), and the converse was also true (*t_18_*=3.03, *p*=.0072). Regarding negative confidence signals, we found significant weights on activity recorded during the Belief task in all but one ROI identified from the Time task (Fig. 4): in right lateral prefrontal cortex (rPFC, *t_18_*= −3.18, *p*=.0052), medial prefrontal area (dACC, *t_18_*=-3.45, *p*=.0029), right parietal cortex (rPAR, *t_18_*= −5.10, *p*=7.55×10^-5^) and inferior parietal lobule (IPL, *t_18_*=-5.56, *p*=2.8×10^-5^), but not in lPFC (*t_18_*=-1.68, *p*=.1095). Conversely, we found significant weights on activity recorded from the Time task in all ROIs identified from the Belief task: SFG (*t_18_*=-4.35, *p*=3.9×10^-5^), MFG (*t_18_*=-2.67, *p*=.0157), rPFC (*t_18_*=-4.90, *p*=1.14× 10^-4^), FPC (*t_18_*=-3.60, *p*=.0020), rPAR (*t*_18_=-3.28, *p*=.0042), and lPAR (*t*_18_=-2.66, *p*=.0160) (Fig. 4). Note however that despite similar labels being used for readability, the extent of each ROI slightly differed between the Belief task, the Time task and the union of tasks (see Supplementary Table 1 for a full description).

**Figure 4.**
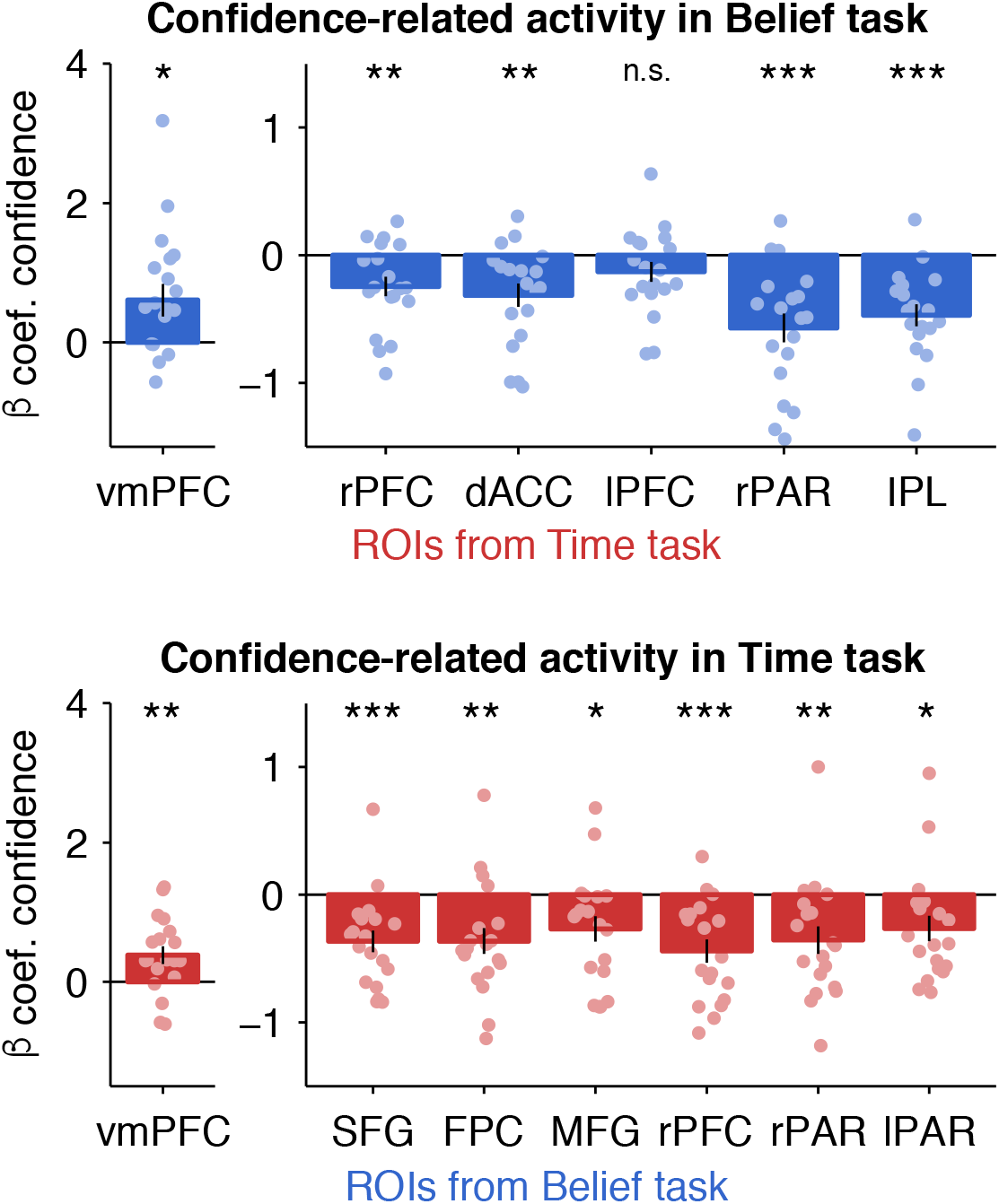
Cross-over analysis of confidence-related activity. Regression coefficients (beta weights) of confidence against activity recorded in one task were extracted from group-level ROIs showing significant association with confidence in the other task. Left and right plots show positive and negative confidence weights on activity recorded in the Time and Belief tasks (red and blue bars). Bars and error bars indicate mean and SEM across participants and dots indicate individual data points (N=19). Abbreviations: vmPFC (ventromedial prefrontal cortex), FPC (frontopolar cortex), dACC (dorsal anterior cingulate cortex and supplementary motor area, PFC (prefrontal cortex), SFG (superior frontal gyrus), MFG (middle frontal gyrus), PAR (parietal cortex), l (left) and r (right). *p<.05, **p<.01, ***p<.001, n.s. not significant, two-tailed paired t-tests. See also Supplementary Table 1.

Finally, to search for any domain-specific activity, we computed two contrasts: confidence in belief judgments minus confidence in time estimates, and vice versa. No brain region showed a stronger association (neither positive or negative) with confidence ratings in one task or the other (all p_FWE_clu_>.424). Note that this absence of significant difference does not allow to conclude that there is no domain-specific confidence computation in the brain. Instead, we can conclude about the existence of a domain-general brain network signaling confidence with increasing activity in some regions (vmPFC) and decreasing activity in others (dorsolateral prefrontal and parietal cortices, plus medial prefrontal regions).

### Confidence signals for the alternative task

Despite the lack of behavioral evidence for a confidence leak from the Belief task to the Time task, it remains possible that confidence in belief judgments may still be represented in the brain, but not translated into a behavioral influence. To search for a correlate of confidence in belief judgments while participants performed the Time task, we added three regressors modulating activity at stimulus onset with behavioral measures taken from the post-scanning Belief task session: response (true/false), accuracy (correct/incorrect), and confidence rating (GLM2, see Methods). We found no confidence-related activity surviving p_FWE_clu_<.05 (for a cluster defining threshold of p=.001 uncorrected, all p_FWE_clu_>.99) (Fig. S3A). We reasoned that despite a lack of neural activity associated with confidence, participants might still compute the true/false response associated with each statement, but we found no cluster reflecting such belief-related activity either (all p_FWE_clu_>.23). We also searched for activity reflecting accuracy (correct/incorrect), an intermediate step in computing confidence, but we again found no significant cluster (all p_FWE_clu_>.43).

For completeness, we searched for confidence leaks using simpler GLMs, with only confidence in belief judgments, either at stimulus onset (Fig. S3B) or at response onset (Fig. S3C). Again, we observed no significant confidence-related cluster (all p_FWE_clu_>.41). Finally, instead of whole-brain analyses, we searched for activity signaling confidence in belief judgments during the Time task in the ROIs identified from the Belief task (ROIs from Fig. 4, see Methods). We found no significant confidence weights (Fig. S3D), neither in the vmPFC (*t_18_*=1.087, *p*=.29) nor in the ROIs negatively associated with confidence: SFG (*t_18_*=-.46, *p*=.65), MFG (*t_18_*=.066, *p*=.95), rPFC (*t_18_*=-.54, *p*=.59), FPC (*t_18_*=-.188, *p*=.85), rPAR (*t_18_*=.013, *p*=.99) and lPAR (*t_18_*=-.70, *p*=.49). In sum, none of these analyses provided any neural evidence for a confidence leak from the Belief task to the Time task.

### Control analysis: neural activity associated with first-order tasks

To conclude about domain-generality of confidence signals, it is important to check that the first-order tasks trigger different cognitive processes (belief judgment and time estimation) and hence involve distinct brain networks. Indeed, direct contrasts between Belief and Time tasks at stimulus onset (GLM1) elicited significantly different activation clusters (Supplementary Table 2), with higher activity in the posterior cingulate cortex (p_FWE_clu_<.001), angular gyrus and mid-occipital cortex (p_FWE_clu_=.005) for belief judgment and in bilateral inferior parietal lobule (both p_FWE_clu_<.001), bilateral primary visual cortex (both p_FWE_clu_<.001) and bilateral inferior orbitofrontal cortex (left: p_FWE_clu_=.004 and right: p_FWE_clu_<.001) for time estimation.

Moreover, the contrast between stimulus categories (history vs. geography statements) yielded significant differences for belief judgment (Supplementary Table 3), but not for time estimation. During the Belief task, we found three separate clusters for geography statements – in bilateral precuneus (p_FWE_clu_<.001), in the inferior and middle temporal gyri (p_FWE_clu_<.001), and in the parahippocampal gyrus and fusiform area (p_FWE_clu_=.041) – while for history statements we found thirteen separate clusters, particularly in the middle temporal cortex (BA21) (p_FWE_clu_<. 001) and posterior cingulate gyrus (p_FWE_clu_<.001) (Supplementary Table 3). In contrast, during the Time task, the contrast between history and geography statements yielded virtually no activation.

Together, these results provide further evidence for largely separate brain networks processing the first-order responses in the two tasks, which contrasts with the largely shared brain network forming confidence judgments in these different first-order responses. In addition, the lack of differential activity between history and geography statements while participants were estimating time further suggests that there was no belief judgment automatically generated during the Time task.

### Exploratory analysis: independent contributions of confidence-signaling regions

Having established a domain-general network for confidence, we further sought to understand whether confidence-related activity in the different ROIs had a significant independent contribution to confidence expressed in the behavior, or whether they simply mirrored each other in a redundant manner. Since the ROIs identified in the union contrast of both tasks (Fig. 3A) depend on statistical thresholding, we pooled all the negative ROIs together (Fig. 5A) and entered their deconvolved, average activity into a multiple regression analysis in which they competed with deconvolved, positive confidence-related activity (averaged over vmPFC voxels) for explaining variance in confidence rating (see Methods).

**Figure 5:**
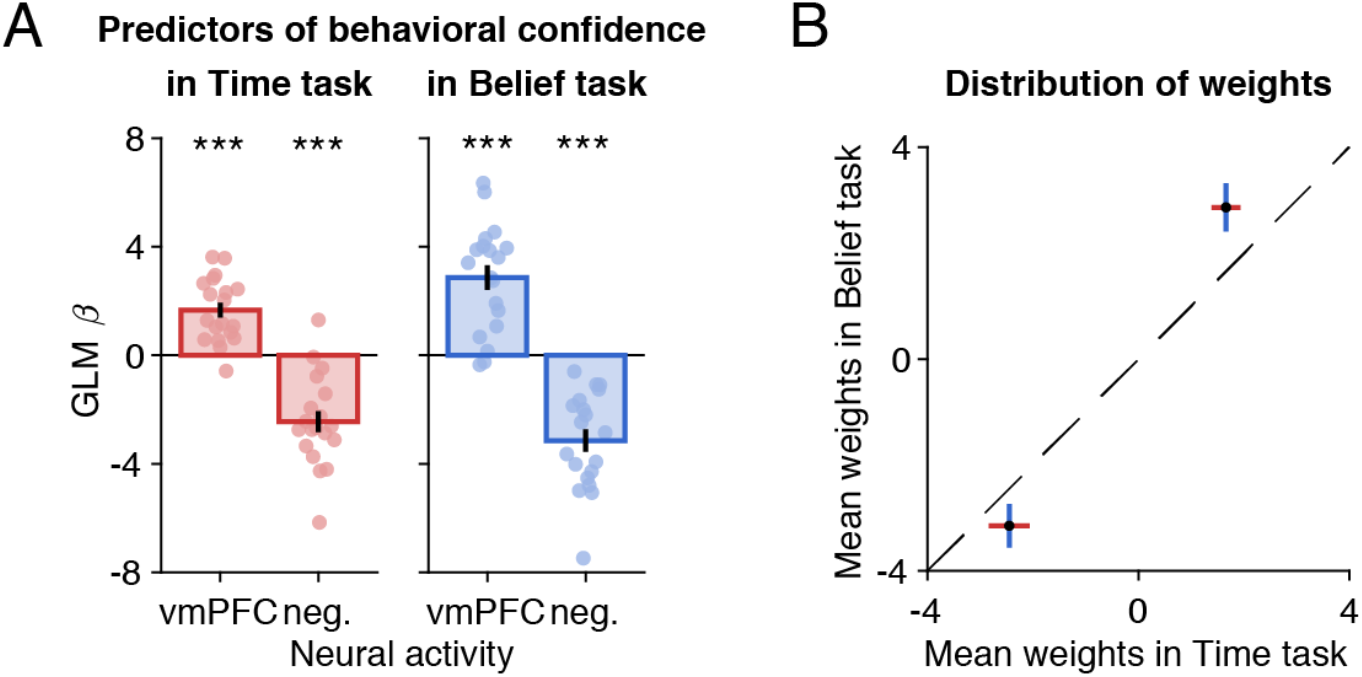
Relative contribution of confidence-related brain activity to behavioral confidence. A) Regression weights of mean neural activity in positive and negative confidence-related regions, competing for variance in behavioral confidence ratings. B) Correlation of weights associated with each ROI between tasks. Note that no group inference was made from these two data points, they just illustrate how weights were positioned with respect to the diagonal. Bars and error bars indicate mean and S.E.M. across participants (N=19). *p<.05, **p<.01, ***p<.001, one-sample t-test against zero for regression coefficients in each ROI. Red: Time task, Blue: Belief task. Abbreviations: ventromedial prefrontal cortex (vmPFC), neg. (negative confidence network).

In both tasks, we found that both the vmPFC (Time: *t_18_*=6.11, *p*=8.9×10^-6^; Belief: *t_18_*=6.32, *p*=5.85×10^-6^) and negative ROIs (Time: *t_18_*=-6.3, *p*=5.8×10^-6^; Belief: *t_18_*=-7.6, *p*=4.97×10^-7^) significantly contributed to confidence expressed in ratings. The positive and negative confidence networks were therefore not redundant, and instead signaled independent information about confidence. This was not true within the negative network: when including the different prefrontal and parietal ROIs as separate regressors in the model, none of them consistently remained a significant predictor in both tasks, as if they were competing for the same variance in confidence ratings.

We also examined the relative weights of positive and negative activity (Fig. 5B): critically, mean weights in both tasks were close to the diagonal (Fig. 5B), indicating that positive and negative networks had a similar relative contribution in the two tasks. For each of the two tasks separately, we found significant anti-correlations between positive and negative activity weights (Time task: ρ=-0.63, *p*=.004; Belief task: ρ=-0.79, *p*=.0001). These results indicate that, within each task, participants who had a larger (more positive) contribution of vmPFC activity to behavioral confidence reports also had a larger (more negative) contribution of the prefronto-parietal network, and the other way around.

Finally, we examined whether the weights of brain activity on confidence reports were correlated between tasks (across participants), separately for the positive and negative networks (Fig. S4). We found borderline correlations for both positive (vmPFC: *ρ*=0.51, *p*=.027) and negative (prefronto-parietal network: *ρ*=0.45, *p*=.054) networks, meaning that participants who had a larger contribution of brain activity to confidence report in one task tended to also have a larger contribution in the other task, as should be the case for a domain-general metacognitive system.

Thus, not only was confidence similarly influenced by the different behavioral factors in the two tasks, but also it was similarly influenced by activity in the different underlying brain regions.

## DISCUSSION

Subjective confidence judgments are pervasively associated with our memories or perceptions and have a major impact on our choices (Fleming et al. 2012; Desender et al. 2018; Rouault, McWilliams, et al. 2018). Here, we leveraged a novel fMRI paradigm to address a fundamental question about how the brain generates confidence judgment: would each domain-dedicated brain system provide a domain-specific confidence signal about its own output, or would a separate shared network compute a domain-general confidence signal based on the output of any domain-dedicated brain system? Our results bring evidence for the second scenario, with confidence judgments being influenced by the same behavioral factors and expressed in the same brain regions during tasks as different as time estimation and belief judgment, classically related to distinct cognitive domains (duration perception vs. semantic memory).

A prerequisite for our demonstration was to compare tasks involving distinct cognitive processes and separate brain systems. We were careful to use the same notion of confidence in both tasks, defined as the subjective probability that a behavior is correct. In both cases, the (first-order) behavioral response was a binary decision, and confidence was expressed in a (second-order) behavioral response – a rating on a visual scale. Thus, we set aside another meaning of confidence as the reliability or precision of representations (Pouget et al. 2016). We also tightly matched the stimuli (history and geography statements) and the movements (left or right button presses) required for responding. What differed, at the cognitive level, was that the information extracted from the stimulus was semantic meaning in one task and time estimation in the other task. Consistently, at the neural level, we found that the two first-order tasks activated anatomically separate brain systems. Moreover, contrary to what was observed during belief judgment, the two stimulus categories (history vs. geography) did not yield distinct activations during time estimation, as if participants were not even processing their semantic content. Together, these findings indicate that we were able to successfully dissociate the first-order tasks, at both the cognitive and neural levels.

Although we observed first-order representations required to solve the two different tasks in separate brain regions, second-order confidence judgments were located in a similar set of brain regions. During both tasks, positive associations with confidence judgments were found in the vmPFC, and negative associations in a large set of brain regions including the medial and lateral prefrontal cortex, the posterior parietal cortex and the frontopolar cortex. The positive association with vmPFC activity is consistent with previous neuroimaging work involving the vmPFC and adjacent perigenual ACC regions in confidence judgments about perceptual responses (Bang and Fleming 2018; Gherman and Philiastides 2018; Rouault and Fleming 2020), about age estimation and likeability rating (Lebreton et al. 2015; Lopez-Persem et al. 2020), about value-based choices (De Martino et al. 2013) and decisions relying on working memory (Morales et al. 2018). The vmPFC therefore appears as a key hub for confidence signals, which suggests that being confident may be intrinsically valuable (Lebreton et al. 2015; Lee and Daunizeau 2021), since the vmPFC is also central to valuation signals generated during both rating and choice tasks (Lebreton et al. 2009; Bartra et al. 2013; Clithero and Rangel 2014; Pessiglione and Daunizeau 2021). We have not tested here whether vmPFC activity was affected by the requirement to provide confidence ratings, but it is likely that similar signals would be generated in the absence of explicit reports, as observed in previous work (Lebreton et al. 2015; Bobadilla-Suarez et al. 2020; Lopez-Persem et al. 2020). One may speculate, given the present and previous results, that a key function of the vmPFC confidence signal may be to promote continuation with the same task or strategy when the behavior is successful (Donoso et al. 2014; Rouault and Fleming 2020). Even if they did not survive our statistical threshold, other regions of the brain valuation system might also contribute to this behavioral regulation function. In particular, we did not find any significant association with confidence in ventral striatum activity, contrary to a recent meta-analysis of metacognitive judgments (Vaccaro and Fleming 2018). It is possible that our sample size did not allow us to reliably identify whether there was any significant confidence-related activity in the ventral striatum, and if so, whether it would be part of a task-general network.

Negative correlation with activity in the bilateral prefronto-parietal network is reminiscent of previous results that associated decreasing confidence to parietal activation during perceptual and memory tasks (Hebart et al. 2015; Bang and Fleming 2018; Vaccaro and Fleming 2018). In addition, SMA and dACC regions have been associated with error detection and conflict monitoring (Holroyd and Coles 2002; Yeung et al. 2004; Shenhav et al. 2013), while the lateral prefrontal and parietal regions form the classic cognitive control network (Koechlin et al. 2003; Owen et al. 2005). Thus, one function of the negative confidence signal may be to recruit cognitive control as is observed with overt errors, for instance to adapt speed-accuracy trade-off or to pay more attention for future decisions (Braun et al. 2017; Desender et al. 2019). Another function may be to drive switching away from the ongoing task or strategy and exploring alternative options (Kolling et al. 2012; Donoso et al. 2014), including seeking information about novel possibilities (Stoll et al. 2016; Rouault et al. 2021). If correct, such considerations would suggest that activity in this prefronto-parietal network arises from the need to monitor and adjust performance, and not from the need to report a confidence rating.

When we contrasted confidence-related activity between tasks, we found no significant activation, hence no evidence for domain-specific confidence signals. This absence of evidence does not imply that there is no domain-specific representation of confidence in the brain. Such domain-specific representations have been previously reported using multivariate analyses based on regions-of-interest (Morales et al. 2018). Nevertheless, our findings do suggest that a substantial part of neural confidence processing is shared between distinct cognitive domains. Our results thus extend a previous study reporting univariate confidence brain signals that were general to a visual memory and a visual perception task (Morales et al. 2018). A shared architecture may be adaptive to centralize the information that guides ongoing behavior, in particular decisions to stay on task or switch away. It is also in line with the notion of a “common currency” for confidence that may allow comparison between tasks (de Gardelle and Mamassian 2014) and hence wise decision about the next course of action. Finally, it may be useful to generalize confidence to new tasks, by providing a proxy for self-ability in situations one has never encountered before.

However, while the contributions of the different brain regions signaling confidence, whether positively or negatively, were similar for belief judgment and time estimation, they were not redundant. Note that the notions of non-redundancy and domain-generality are orthogonal: the positive and negative confidence brain networks may play two different roles, but still each play the same role whatever the cognitive processes engaged to solve the task. The non-redundancy is an important clue for the functional roles of the two networks, because certain brain systems are typically observed to activate in opposition to one another, so it could have been the case that negative regions simply mirror the information content of positive regions. Instead, the current findings suggest that the two networks may integrate separate, independent aspects of confidence that future studies should seek to tease apart. A first functional interpretation is that positive activity in a valuation region such as the vmPFC, engaging motivational circuitry, reflects a preference for high-confidence situations, while negative activity in the prefronto-parietal network signals cognitive control involvement to cope with difficult situations in which confidence is too low. Another functional interpretation is that the vmPFC monitors ongoing beliefs about performance at the task level (‘global’ confidence), while the prefronto-parietal network monitors trial-by-trial (‘local’) confidence at the decision level (Wittmann et al. 2016; Rouault and Fleming 2020). A third possible functional interpretation is that the prefronto-parietal network estimates decision uncertainty as a computational variable, while the vmPFC generates the associated subjective experience, for instance a positive feeling associated with high confidence (D’Argembeau 2013; Bang and Fleming 2018).

At the behavioral level, the factors known to influence confidence (in particular accuracy, RT and difficulty) had similar weights in the belief judgment and time estimation tasks. Although a regression is not a mechanistic model, these results suggest that confidence in the two tasks was computed from the same building blocks. They are consistent with a previous study that compared perceptual and knowledge-based decision tasks and showed that ratings in both cases complied with normative expectations from a statistical confidence model (Sanders et al. 2016). In addition, there was a borderline correlation across participants in calibration and discrimination metrics that describe how confidence varies with performance. Calibration measures indicated that most participants were overconfident in their performance, in line with previous observations (Berner and Graber 2008; Moore and Healy 2008; Lebreton et al. 2019). Such correlations in metacognitive ability metrics have been viewed as evidence for domain-generality of confidence, the rationale being that if the computation of confidence is domain-general, participants who are good at discriminating their own high versus low performance should be so in any domain (for a review, see (Rouault, McWilliams, et al. 2018). However, previous studies relying on such correlations have provided mixed results as to whether metacognitive ability correlates across tasks (McCurdy et al. 2013; Faivre et al. 2017; Samaha and Postle 2017; Mazancieux et al. 2020) or not (Baird et al. 2013; Valk et al. 2016; Fitzgerald et al. 2017). Nevertheless, correlation approaches may only provide indirect evidence, as a positive correlation could be driven by a third factor (such as compliance with instructions), while nonsignificance does not prove an absence of correlation and might be due to other reasons (see (Shekhar and Rahnev 2020). In addition, a positive correlation at the behavioral level can coexist with anatomical and functional separation between underlying metacognitive systems at the neural level (Baird et al. 2013; McCurdy et al. 2013). Here, the conclusions that could be drawn from interparticipant correlations are even more limited due to the small sample (N=20). In future studies, it would be interesting to examine whether and how individual traits impact behavioral confidence reports and neural confidence signals, using larger (Rouault, Seow, et al. 2018) and appropriately powered sample sizes.

We observed no evidence for a leak between tasks, in that confidence in belief judgment was not observed at the neural level during the time estimation task, and did not affect at the behavioral level confidence in decisions about durations. We initially reasoned that the presence of a confidence leak could have been an indirect marker of domain-generality: even if task-specific metacognitive systems were partially dissociable, a leak would denote the existence of a metacognitive representation that integrates the information related to different tasks, hence forming a domain-general confidence signal. It should be noted, however, that an absence of confidence leak is no proof of domain-specificity, since it could be the signature of a unique but efficient domain-general metacognitive system that would be immune to misattribution (i.e., that would track with which task confidence representations are associated with fidelity). Besides, this null result should not be taken as evidence against the possibility of a confidence leak, as it may come from limitations in our design or measure. First, confidence in belief judgment was quite low on average, as the mean accuracy was 67% correct. Therefore, the uncertainty level may remain too high to influence confidence in other responses, if confidence is to leak only when one is sure enough. Second, a confidence leak in our design required the first-order response to the alternative task to be automatically generated. This was unlikely to be the case for belief judgment during time estimations, as there was no neural signature of semantic processing. As tasks were blocked, participants knew before seeing the stimulus that they were asked to focus on time, so they might have avoided reading the statement. This situation is therefore critically different from previous paradigms in which confidence leak was observed, where participants were asked to provide two binary responses and related confidence ratings for each stimulus (Rahnev et al. 2015). The absence of a confidence leak when the alternative task is not performed could therefore be interpreted as restricting the property of automaticity, observed for confidence signaling (Lebreton et al. 2015), to situations where a behavioral response is overtly provided. Alternatively, confidence signals could be automatically generated even for covert responses, but appropriately tagged to avoid cross-contamination between tasks and drive behavior in a more adaptive manner.

To conclude, we have compared confidence behavioral ratings and neural signals between tasks that required very distinct cognitive processes. We found that confidence was related to the same behavioral factors and the same neural activations in the two tasks, without creating interferences between tasks. These results suggest the existence of a domain-general neural architecture estimating whether the behavior is correct, which may drive subsequent decisions in an adaptive manner. Our findings do not preclude, however, the existence of other domain-specific neural signals associated with notions of confidence defined as the reliability or precision of mental representations. Further research is needed to investigate these potential domain-specific signals and examine how they articulate with a centralized representation of confidence to drive our thoughts and actions.

## ACKNOWLEDGMENTS

The study was funded by a Starting Grant (BioMotiv, 260747) from the European Research Council. This work has also received support under the program «Investissements d,Avenir» launched by the French Government and implemented by the Agence Nationale de la Recherche (ANR-10-IAIHU-06). The funders had no role in study design or manuscript preparation. M.R. is the beneficiary of a postdoctoral fellowship from the AXA Research Fund. During the time of the research M.L. was a PhD candidate in the MBB lab, supported by a PhD fellowship from the French Ministry of Research. During the preparation of the manuscript, M.L. has been supported by an SNSF Ambizione grant (PZ00P3_174127).

The authors thank Stefano Palminteri and the CENIR team for their help in the fMRI data collection. The authors additionally thank Sebastien Massoni, Jean-Christophe Vergnaud and Guillaume Hollard for helpful discussions about belief and confidence elicitation.

## AUTHOR CONTRIBUTIONS

Conceptualization: M.L. and M.P.; Data Collection: M.L.; Formal Analysis: M.R. and M.L.; Investigation: M.R., M.L. and M.P.; Writing – Original Draft: M.R.; Writing – Review & Editing: M.L. and M.P.; Supervision: M.P.; Funding Acquisition: M.P.

## CONFLICTS OF INTEREST

The authors declare no competing interests.

## Notes

### Competing Interest Statement

The authors have declared no competing interest.

