## Supplementary Material for "A shared brain system forming confidence judgment across cognitive domains"

**ORCIDs :**

Marion Rouault: 0000-0001-6586-3788

Maël Lebreton: 0000-0002-2071-4890

Mathias Pessiglione: 0000-0002-6992-3677

**Supplementary Results**

*Incentivised Belief judgment task ("IncBelief").* We analyzed participants' behavior on the IncBelief task. Comparisons with the Belief task were restricted to the three post-scanner Belief task sessions, such that in both cases participants were seeing the exact same statements for the second time (Fig. 1A). First, we found that participants reached 68.9% correct, an accuracy similar to that of the Belief task ($t_{19}$=1.32, $p$=.202). Participants' mean confidence was slightly higher on the IncBelief than on the Belief task ($t_{19}$=2.40, $p$=.027). This slightly higher confidence under incentivisation did not translate into meaningful differences in metacognitive abilities, as there was no difference in calibration ($t_{19}$=-1.47, $p$=.159) or discrimination ($t_{19}$=-1.37, $p$=.19) between the Belief and IncBelief tasks. In line with the Belief task data from the scanner sessions (Fig. 2A), we found that trial-by-trial confidence ratings were predicted by fluctuations in accuracy ($t_{19}$=5.99, $p$=9.14×10$^{-6}$), RTs ($t_{19}$=-7.89, $p$=2.02×10$^{-7}$), difficulty ($t_{19}$= 8.63, $p$=5.31×10$^{-8}$) and confidence from the previous trial ($t_{19}$=2.69, $p$=.015), all standard predictors of confidence judgments (Fig. S2). We also observed that these results were robust to removing of the regressors of no interest and again found a significant influence of accuracy ($t_{19}$=6.17, $p$=6.2×10$^{-6}$), RTs ($t_{19}$=-7.58, $p$=3.7×10$^{-7}$), difficulty ($t_{19}$=9.15, $p$=2.2×10$^{-8}$), and confidence from the previous trial ($t_{19}$=2.75, $p$=.0127) on confidence ratings. Furthermore, to evaluate the similarity in how these various factors impacted confidence rating, we computed the correlation of their regression weights between Belief and IncBelief tasks. This correlation was strongly significant at the group level (mean ρ=0.60, $t_{19}$=10.2, $p$=3.8×10$^{-9}$). Together, these results provide evidence that the incentivization schedule (IncBelief) on confidence judgments as compared to spontaneous confidence ratings (Belief) did not substantially alter confidence or metacognition on this task, in line with previous reports (Schlag et al., 2015; Hollard et al., 2016).

*Inter-individual correlations.* To assess domain-generality, many studies have exploited inter-individual variations in metacognitive ability, that is the strength of the correspondence between subjective confidence and objective accuracy (Rouault et al., 2018; Hoven et al., 2019). The reasoning is that if metacognition is a domain-general resource, then participants who are good at discriminating their own high versus low performance should be so in any task. We computed several metrics of metacognitive ability, which specify how participants provided confidence ratings in relation to accuracy (see Methods). Because our sample was limited (N=20), we provide Bayes factors (BF) in addition to *p*-values, to give an idea of the strength of the evidence conveyed by null results.

First, we found that confidence level, i.e. the individual tendency to rate confidence high or low on average, did not correlate between tasks ($\rho$=.25, *p*=.28, BF=.814). There was a significant difference in confidence level between Belief and Time tasks ($t_{19}$=-4.43, *p*=2.85×$10^{-4}$, BF=109.8), presumably due to a difference in accuracy between tasks ($t_{19}$=-3.70, *p*=.0015, BF=25.57). In contrast, calibration, i.e., the difference between mean objective accuracy and mean subjective confidence, correlated across participants ($\rho$=.48, *p*=.031, BF=4.772), a trend result that disappeared after removing an outlier participant (*p*=.21) (Fig. S1A). There was also no significant difference between tasks ($t_{19}$=.86, *p*=.39, BF=.323). This finding remains inconclusive in terms of a true correlation in confidence level between tasks, once "corrected" for the difference in accuracy. Finally, we computed discrimination, a metrics of metacognitive ability reflecting how confidence rating differentiates correct from incorrect responses on a trial-by-trial basis (difference in confidence rating between correct and incorrect trials). We found an inconclusive correlation for discrimination ($\rho$=.39, *p*=.086, BF=2.080) (Fig. S1B), which together with calibration provide mixed evidence as to whether metacognitive ability manifested similarly in the two tasks. We note that differences in first-order accuracy between tasks precluded the application of signal detection theory frameworks, such as meta-*d'*, for estimating metacognitive abilities.

**Supplementary Figures and Tables**

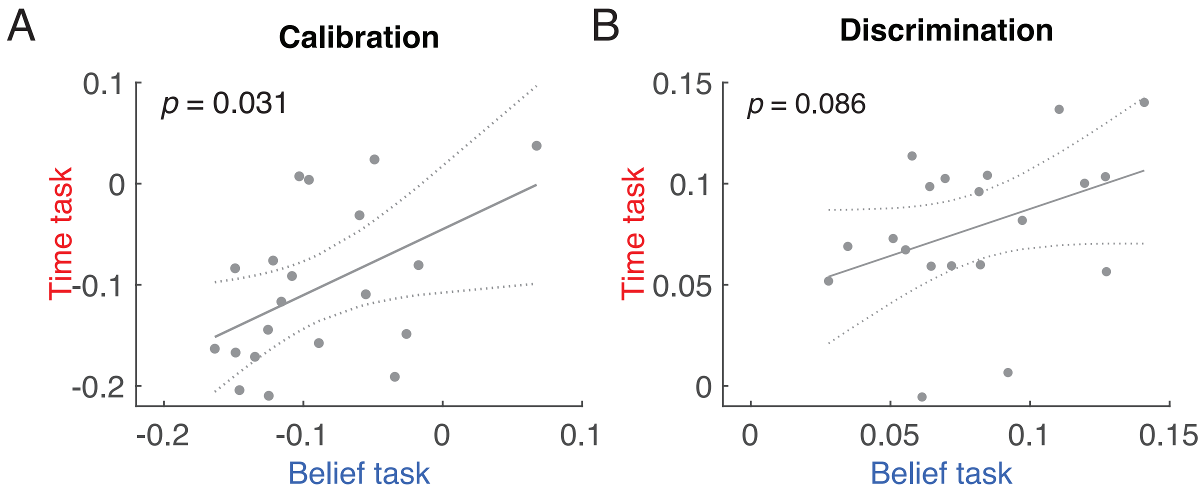


***Supplementary Figure 1. Inter-individual correlations of metacognitive ability measures.***
*Calibration (left panel) is an index of over- or under-confidence (difference between mean confidence and mean accuracy). Discrimination (right panel) is an index of how well confidence tracks variations in accuracy across trials (mean confidence for correct minus mean confidence for incorrect trials). Circles indicate individual participants (N=20). Dotted lines indicate the 95% confidence bounds on the predicted values from a linear regression fit. P-values indicate significance levels of Pearson correlation coefficients.*

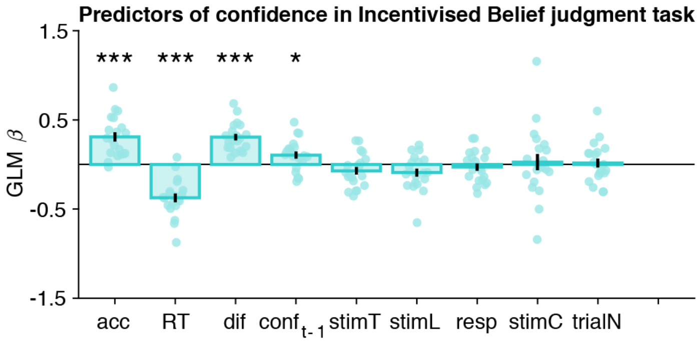


***Supplementary Figure 2. Regression weights of behavioral factors on confidence in the Incentivised Belief judgment task (IncBelief).***

*Behavioral predictors of confidence rating at trial t on the IncBelief task were accuracy (acc), response times (RT) and difficulty level (dif). Regressors of no-interest were included for completeness: confidence at the previous trial ($conf_{t-1}$), stimulus presentation time (stimT), stimulus length (stimL), first-order response (resp, left or right), stimulus category (stimC, historic or geographic), and trial number (trialN). Error bars indicate S.E.M. across participants. Dots indicate individual participants (N=20). *p<.05, ***p<.001, one sample t-tests against zero.*

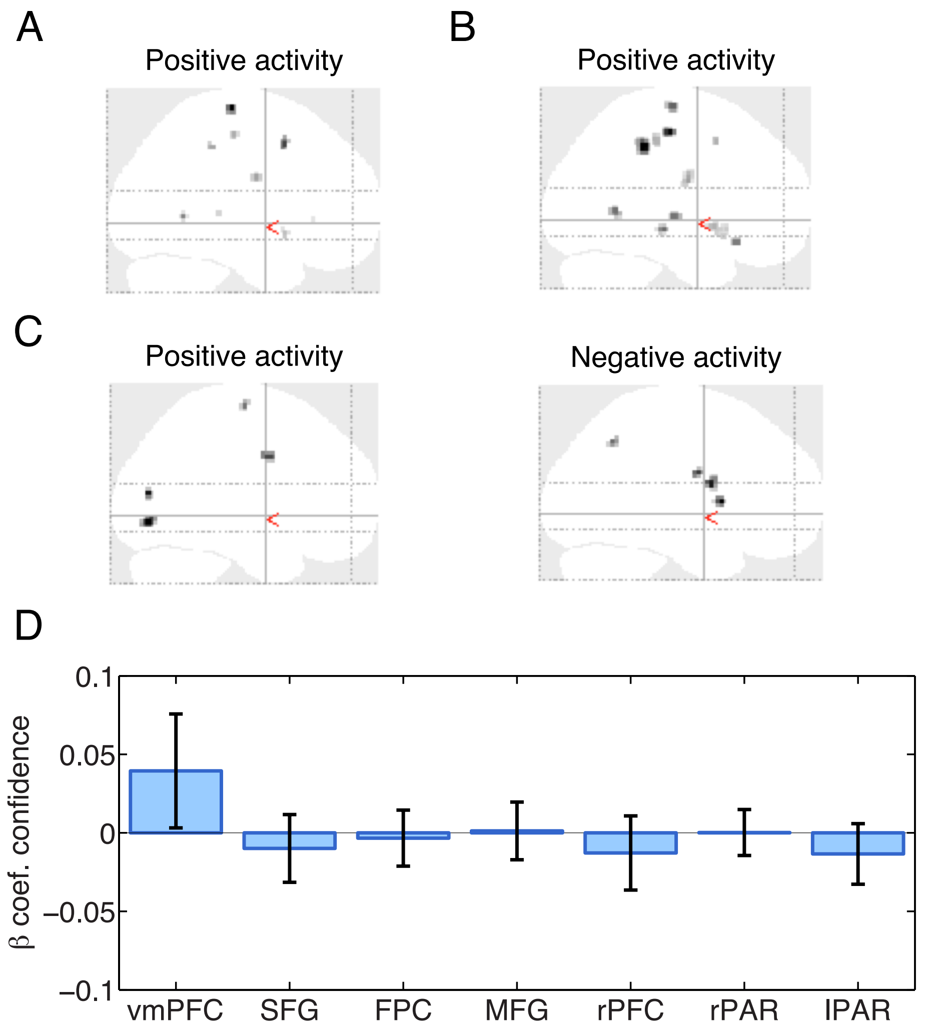


***Supplementary Figure 3: Testing automatic signaling of confidence in the alternative task.***
*A) During the Time task, in a whole-brain GLM, we searched for activity associated with confidence in the Belief task about the same stimuli. Glass brain maps show no positive or negative activity surviving p<.05 cluster-corrected, for a cluster-defining threshold of p=.001 uncorrected. B,C) Similar results were obtained in a GLM with only confidence in the Belief task as an additional parametric modulator (B), and at the response onset instead of at the stimulus onset (C). D) ROI analysis. No significant confidence-related activity for the Belief task during the Time task was found in the ROIs identified from the Belief task scanning sessions. Abbreviations: vmPFC (ventromedial prefrontal cortex), SFG (superior frontal gyrus), FPC (frontopolar cortex), MFG (middle frontal gyrus), PFC (prefrontal cortex), PAR (parietal cortex), l (left) and r (right). Error bars represent SEM across participants (N=19).*

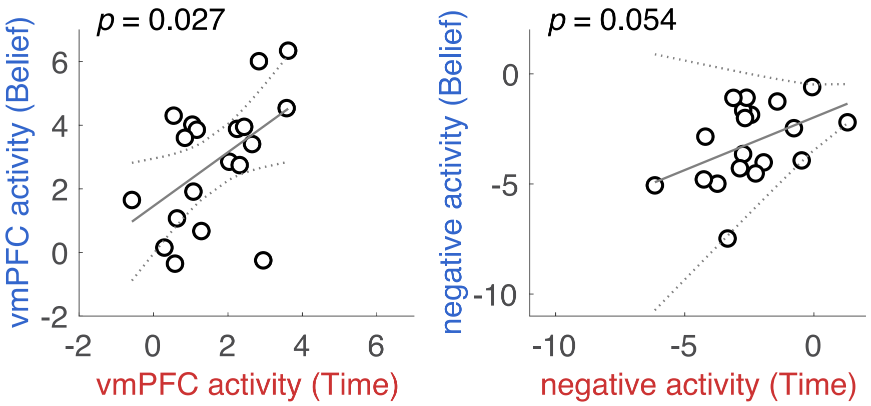


***Supplementary Figure 4.*** *Individual weights (from main Figure 5) of vmPFC activity (left panel) and prefronto-parietal activity (right panel) in the Belief task as a function the same weights in the Time task. p-values indicate statistical significance of the correlations between tasks across participants (N=19).*

| Regions | Later. | pFWE-corr cluster-level | Peak MNI x | y | z | T (peak) | Size (voxels) |
|---|---|---|---|---|---|---|---|
| ***Positive effect of confidence in Belief task*** | | | | | | | |
| Occipital cortex | R | $5.7\times10^{-13}$ | 16 | -76 | -4 | +6.55 | 1572 |
| Precuneus and posterior cingulate cortex | L | 0.0027 | -8 | -50 | 6 | +6.19 | 237 |
| ***Negative effect of confidence in Belief task*** | | | | | | | |
| Inferior parietal lobule and superior parietal cortex (“rPAR”) | R | $<1\times10^{-13}$ | 8 | -60 | 54 | -10.07 | 3557 |
| Inferior parietal lobule (“IPL”) | L | $5.3\times10^{-7}$ | -46 | -54 | 50 | -7.45 | 655 |
| Occipital cortex | N/A | $1.4\times10^{-6}$ | -10 | -78 | -8 | -7.44 | 599 |
| Superior and middle frontal gyrus (“SFG”) | R | 0.0441 | 14 | 38 | 50 | -7.19 | 133 |
| Frontopolar cortex and superior frontal gyrus (“FPC”) | R | $2.4\times10^{-5}$ | 22 | 52 | 38 | -6.17 | 452 |
| Middle frontal gyrus (“MFG”) | R | 0.0031 | 32 | 32 | 38 | -6.10 | 231 |
| Superior frontal gyrus (“rPFC”) | R | 0.0294 | 10 | 10 | 68 | -5.70 | 147 |
| ***Positive effect of confidence in Time task*** | | | | | | | |
| Occipital cortex | R | $2.3\times10^{-13}$ | 12 | -82 | 2 | +8.60 | 1840 |
| ***Negative effect of confidence in Time task*** | | | | | | | |
| MFG/SFG/IFG (BA9/8) (“rPFC”) | N/A | $3.3\times10^{-4}$ | 42 | 14 | 44 | -6.29 | 382 |
| dACC/SMA/preSMA/SFG (“dACC”) | N/A | $2.9\times10^{-10}$ | 12 | 42 | 46 | -6.08 | 1257 |
| MFG/IFG (BA9/8) (“lPFC”) | L | $1.1\times10^{-8}$ | -34 | 22 | 36 | -5.98 | 992 |
| AG/Supramarginal gyrus/Inferior Parietal Lobule (“IPL”) | R | 0.0010 | 48 | -56 | 34 | -5.65 | 308 |
| Occipital cortex | L | 0.0449 | -8 | -84 | 0 | -5.12 | 145 |
| Superior parietal cortex (“rPAR”) | R | 0.0081 | 12 | -68 | 62 | -4.72 | 214 |
| ***Positive effect of confidence pooled across Belief and Time tasks (union)*** | | | | | | | |
| Ventromedial prefrontal cortex | N/A | 0.0042 | -2 | 50 | -14 | +5.76 | 201 |
| Occipital cortex | R | $<1\times10^{-13}$ | 12 | -82 | 4 | +9.49 | 3926 |
| ***Negative effect of confidence pooled across Belief and Time tasks (union)*** | | | | | | | |
| Precuneus and parietal cortex (“rPAR”) | R | $<1\times10^{-13}$ | 52 | -58 | 36 | -9.75 | 4617 |
| Middle and inferior frontal gyrus (“lMFG”) | L | $5.6\times10^{-7}$ | -38 | 18 | 42 | -8.60 | 592 |
| Inferior parietal cortex and superior parietal lobule (“lPAR”) | L | $7.5\times10^{-12}$ | -46 | -44 | 44 | -8.19 | 1250 |
| Dorsal anterior cingulate cortex, supplementary and pre-supplementary motor area, and superior frontal gyrus (“dACC”) | R | $<1\times10^{-13}$ | 26 | 46 | 24 | -8.13 | 3848 |
| Occipital cortex | L | $1.3\times10^{-6}$ | -10 | -76 | -6 | -7.22 | 551 |
| Inferior frontal gyrus (“rIFG”) | R | 0.0070 | 46 | 16 | 14 | -7.07 | 183 |
| Inferior frontal gyrus (“lIFG”) | L | 0.0205 | -50 | 22 | 14 | -5.63 | 147 |
| Middle temporal and inferior temporal gyrus | R | 0.0017 | 54 | -52 | -8 | -5.58 | 234 |
| Frontopolar cortex and superior frontal gyrus (“FPC”) | L | 0.0039 | -24 | 52 | 28 | -5.44 | 204 |

***Supplementary Table 1****: Regions with significant whole-brain confidence-related activity thresholded at $p<.05$ FWE cluster-corrected at a cluster-defining threshold of $p=.001$ uncorrected, $k>10$ voxels, for each contrast computed from GLM1 (see Methods). Later., laterality; L, left; R, right. Cerebellar activations were not analyed and are not reported here. Note that the contrasts regarding each task separately correspond to twice fewer data (a half-session per participant, N=19 participants). Abbreviations are provided in quotes for the areas further analyzed in Figures 3, 4 and 5.*

| Regions | Later. | pFWE-corr cluster-level | Peak MNI x | y | z | T (peak) | Size (voxels) |
|---|---|---|---|---|---|---|---|
| *Contrast Belief minus Time at onset of first-order response* | | | | | | | |
| Posterior cingulate cortex | R | <0.001 | 10 | -50 | 4 | +7.65 | 1159 |
| Angular gyrus and mid-occipital cortex | L | 0.005 | 44 | -74 | 32 | +5.89 | 219 |
| *Contrast Time minus Belief at first-order response* | | | | | | | |
| Cingulate cortex | N/A | <.001 | 0 | -26 | 26 | +11.6 | 3853 |
| Inferior parietal lobule | R | <.001 | 64 | -38 | 34 | +9.74 | 3432 |
| Inferior parietal lobule | L | <.001 | -52 | -60 | 48 | +8.29 | 1125 |
| Occipital cortex and lingual gyrus | L | <.001 | -12 | -92 | -2 | +7.51 | 653 |
| Superior and middle frontal gyrus | L | 0.002 | -30 | 54 | -10 | +7.03 | 249 |
| Inferior frontal and precentral gyrus | R | <.001 | 42 | 4 | 30 | +7.00 | 537 |
| Superior frontal gyrus | R | 0.019 | 14 | 52 | 38 | +6.93 | 166 |
| Inferior orbitofrontal cortex | L | 0.004 | -18 | 10 | -24 | +6.90 | 229 |
| Inferior orbitofrontal cortex and middle frontal gyrus | R | <.001 | 42 | 38 | -4 | +6.76 | 511 |
| Frontopolar cortex and superior frontal gyrus | R | 0.050 | 18 | 64 | -6 | +6.62 | 131 |
| Medial and inferior frontal gyrus | R | 0.009 | 12 | 12 | -16 | +6.12 | 194 |
| Insula and inferior frontal gyrus | R | 0.001 | 36 | 14 | -2 | +6.11 | 276 |
| Occipital cortex and lingual gyrus | R | <.001 | 16 | -100 | 12 | +5.80 | 422 |

***Supplementary Table 2: Neural correlates of first-order tasks reveal a large domain-specificity***
*Regions with significant whole-brain confidence-related activity thresholded at p<.05 FWE cluster-corrected at a cluster-defining threshold of p=.001 uncorrected, k>10 voxels, for each contrast at stimulus onset (see Methods). Later., laterality; L, left; R, right. BA: Brodmann area. N=19 participants.*

| Regions | Later. | pFWE-corr cluster-level | Peak MNI x | y | z | T (peak) | Size (voxels) |
|---|---|---|---|---|---|---|---|
| ***Contrast History minus Geography in Belief task at onset of first-order response*** | | | | | | | |
| Middle temporal cortex | L | <.001 | -64 | -44 | -2 | 12.26 | 1970 |
| Precuneus and posterior cingulate gyrus | N/A | <.001 | -2 | -44 | 30 | 9.71 | 1036 |
| Anterior cingulate cortex | N/A | <.001 | -6 | 56 | -10 | 9.21 | 621 |
| Angular gyrus and inferior parietal lobule | L | <.001 | -42 | -70 | 36 | 8.34 | 734 |
| Middle temporal gyrus | R | <.001 | 50 | -6 | -18 | 7.83 | 632 |
| Angular gyrus and inferior parietal lobule | R | <.001 | 46 | -72 | 42 | 7.29 | 620 |
| Middle frontal gyrus | L | .001 | -40 | 12 | 58 | 6.98 | 253 |
| Frontopolar cortex | R | <.001 | 10 | 64 | 24 | 6.59 | 513 |
| Superior frontal gyrus | L | <.001 | -20 | 36 | 44 | 6.29 | 387 |
| Inferior occipital gyrus | L | .021 | -38 | -78 | -10 | 5.93 | 152 |
| Middle temporal gyrus | R | .008 | 66 | -42 | -2 | 5.80 | 186 |
| Middle occipital gyrus | R | <.001 | 6 | -86 | -6 | 5.21 | 353 |
| Superior frontal gyrus | R | .020 | 26 | 30 | 50 | 5.14 | 153 |
| ***Contrast Geography minus History in Belief task at onset of first-order response*** | | | | | | | |
| Bilateral precuneus | L | <.001 | -18 | -62 | 16 | 11.89 | 5012 |
| Inferior and middle temporal gyrus | L | <.001 | -56 | -62 | -8 | 9.26 | 301 |
| Parahippocampal gyrus and fusiform area | L | .041 | -28 | -42 | 12 | 6.90 | 129 |

***Supplementary Table 3: Neural correlates of history vs. geography statements in the first-order Belief task***

*Regions with significant whole-brain confidence-related activity thresholded at p<.05 FWE cluster-corrected, at a cluster-defining threshold of p=.001 uncorrected, k>10 voxels, for each contrast at stimulus onset (see Methods). Cerebellar activations were not analyzed and are not reported here. Later., laterality; L, left; R, right. BA: Brodmann area. N=19 participants.*
